# Piezo1 transduces mechanical signals to inhibit osteoclast fusion and coordinate bone homeostasis

**DOI:** 10.64898/2026.06.02.727841

**Authors:** Yao Wang, Han Wang, Tian you Kan, Junqi Cui, Xuran Li, Kai Yuan, Liao Wang, Mengning Yan, Linyang Chu, Hanjun Li, Zhifeng Yu

## Abstract

Osteoclasts originate from bone marrow–derived macrophages, and their maturation relies on fusion of pre-osteoclasts into mature osteoclasts that maintain bone homeostasis. Previous studies established that mechanical signals regulate osteoblast activity in bone homeostasis, although whether such signals regulate bone homeostasis by acting on osteoclasts remains unclear. Herein, membrane tension decreased during pre-osteoclast fusion, accompanied by reduced Piezo1. This suggests that mechanical stimulation inhibits fusion of monocytes into osteoclasts through Piezo1. Piezo1-knockout monocytes (*CTSK^Cre^; Piezo1^fl/fl^*) attenuated the anabolic effect of exercise on bone mass in vivo, whereas shear stress inhibited osteoclast fusion in vitro. Piezo1 activation elevated E-cadherin, which anchors Merlin. This led to hyperactivation of the Yes-associated protein (YAP) signaling pathway, subsequently activating TAX1BP1 and suppressing NF-κB signaling as well as osteoclast differentiation and maturation. Thus, physical exercise activates a Piezo1–E-cadherin–Merlin–YAP axis that prevents osteoclast fusion, presenting a druggable mechanotransduction pathway for osteoporosis.

## INTRODUCTION

During the fusion of monocytes/osteoclasts, the cells undergo significant morphological and volumetric changes accompanied by cytoskeletal rearrangement ^1^. These changes induce structural deformation and alter the curvature of the cell membrane lipid bilayer, which are key sources of membrane tension ^2^. Specifically, osteoclast precursor cells initially recognize each other through surface molecules (e.g., DC-STAMP) ^3^. Subsequently, they form adhesive junctions through surface proteins (e.g., E-cadherin), bringing the cell membranes of adjacent cells into close contact and inducing cytoskeletal remodeling ^4^. Finally, under the action of fusion proteins (e.g., DC-STAMP), they complete cell fusion and volume expansion ^3,5^. Studies have shown that membrane fusion alters membrane tension, which in turn affects membrane fusion ^6,7^. We hypothesize that the membrane tension of osteoclasts may undergo corresponding changes during fusion.

Studies have shown that the conformation of Piezo1 is sensitive to subtle variations in membrane tension, such as structural deformation and curvature in the lipid membrane ^8^. Thus, changes in the membrane tension can directly activate the Piezo1 signaling pathway. As a mechanosensitive cation channel, Piezo1 opens upon mechanical stimulation and directly mediates calcium ion influx, converting physical signals into intracellular biochemical signals ^9,10^. Calcium ions are essential signaling molecules for osteoclast maturation; they activate calcineurin to induce the dephosphorylation and activation of nuclear factor of activated T cells c1 (NFATc1)—a key transcription factor for osteoclastogenesis that directly regulates the expression of osteoclast-specific marker genes such as tartrate-resistant acid phosphatase (*TRAP*) and cathepsin K (*CTSK*) ^11^. Furthermore, cryo-electron microscopy has confirmed that Piezo1 is structurally associated with E-cadherin ^12^, which facilitates cell–cell recognition and aggregation and promotes osteoclast fusion ^13^. However, whether mechanics influence osteoclast fate through Piezo1 remains unknown.

Herein, we investigated changes in membrane tension during osteoclast fusion and examined the role of Piezo1 in osteoclast differentiation. We hypothesized that mechanical stimulation is not required for the fusion of monocytes into osteoclasts and that the application of mechanical stimulation or activation of Piezo1 would inhibit osteoclast maturation and thereby regulate bone metabolism.

## RESULTS

### Osteoclast-specific Piezo1 knockout attenuates the anabolic effect of treadmill exercise on bone mass

We crossed *CTSK ^Cre^* mice with *Piezo1 ^fl/fl^* mice to generate *CTSK ^Cre^; Piezo1 ^fl/fl^* mice. Subsequently, we measured the bone mass of wild-type (WT) mice and *CTSK ^Cre^*; *Piezo1 ^fl/fl^* mice (CTSK-P1) under two conditions: sedentary and exercise (Figure 1. A). Osteoclast-specific Piezo1 knockout did not affect bone mass in mice under sedentary conditions, which is consistent with previous studies ^14^. After subjecting the mice to 8 weeks of running exercise, the exercise regimen significantly increased bone mass in the mice (BV/TV, from an average of 6.2% to 13.5%) (Figure 1. B–C). Based on previous studies, this phenomenon can likely be attributed to osteoblasts amplifying the effect of mechanical stimulation on bone mass ^14,15^. However, surprisingly, the increase in bone mass observed in CTSK-P1 mice was significantly lower than that in WT mice. In other words, osteoclast-specific Piezo1 knockout attenuated the positive effect of running on bone mass in mice (BV/TV, from an average of 13.5% to 11.1%). Thus, osteoclasts may also be one of the target cells through which mechanical stimulation regulates bone mass.

**Figure 1.**
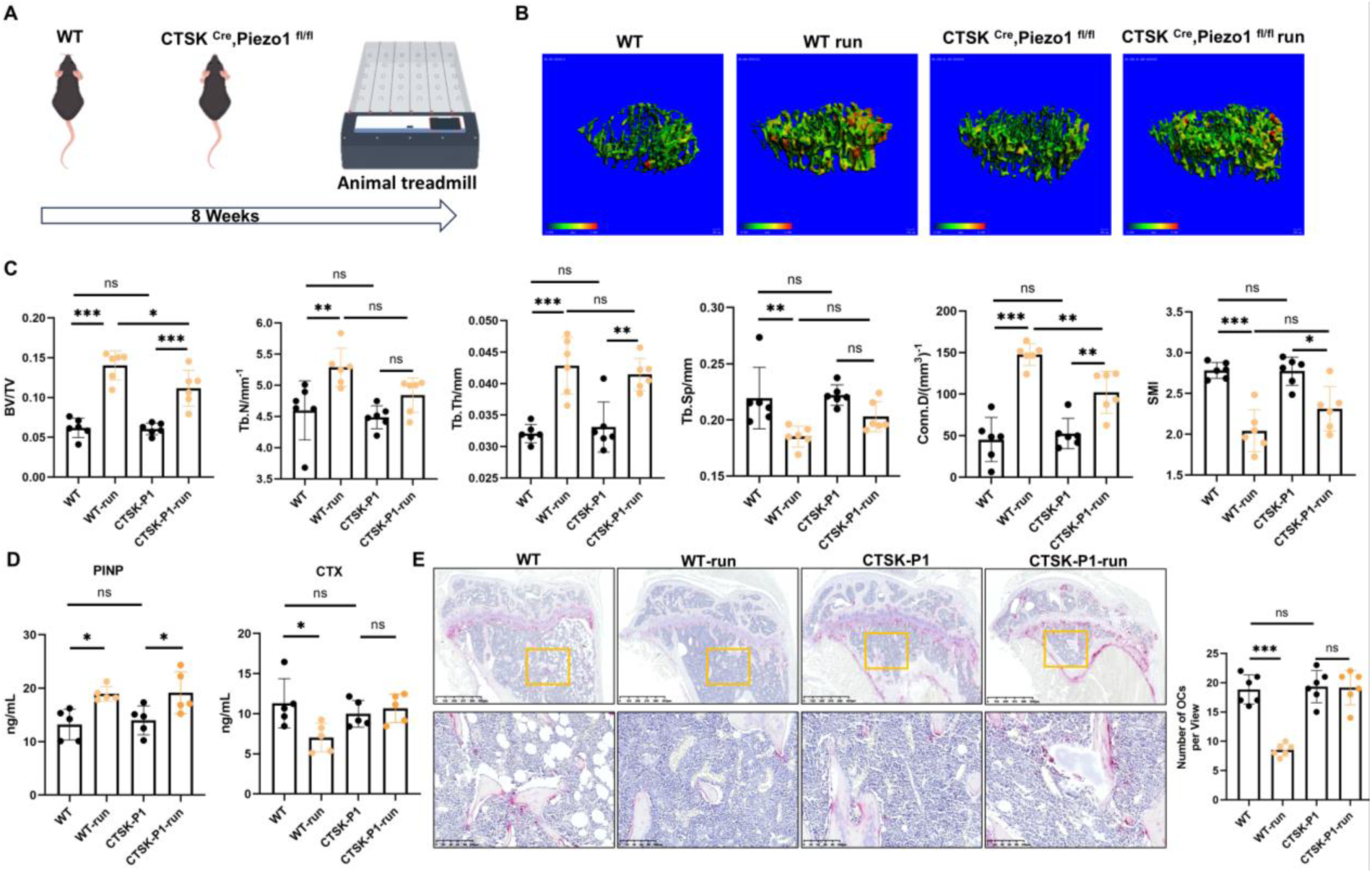
Osteoclast-specific Piezo1 knockout attenuates the promotion of running exercise on bone mass. **(A)** Schematic of mouse running protocol. **(B)** 3D reconstruction of wild-type mice and *CTSK ^Cre^; Piezo1 ^fl/fl^*Mice via micro-CT. **(C)** Parameter statistics of tibial trabeculae, including BV/TV, Tb.N, Tb.Th, Tb.Sp, Conn.D, and SMI (*n*=6). **(D)** Enzyme-Linked Immunosorbent Assay (ELISA) detection of serum CTX and PINP (*n*=6). **(E)** TRAP staining was conducted on bone tissue sections, and osteoclast quantification was performed subsequently (*n*=6). Data are expressed as the mean ± SD. * *P*<0.05, * * *P*<0.01, *** *P*<0.001.

Furthermore, treadmill exercise increased the level of procollagen type I N-propeptide (PINP) in the serum of mice while decreasing that of C-terminal telopeptide (CTX) (Figure 1. D). This further suggests that both osteoblasts and osteoclasts are affected by mechanical stimulation. Tissue sections confirmed that treadmill exercise reduced the number (TRAP staining) of osteoclasts (Figure 1. E). Similarly, under sedentary conditions, osteoclast-specific Piezo1 knockout did not affect the serological and histological phenotypes of mice.

In summary, we identified osteoclasts as novel target cells for the regulation of bone mass by mechanical stimulation, which challenges the traditional notion that mechanical stimulation modulates bone mass solely through osteoblasts. Mechanical stimulation (treadmill exercise) can bidirectionally regulate the functional activity of osteoblasts and osteoclasts. Furthermore, Piezo1 in osteoclasts exhibits strict conditional specificity in regulating bone mass: the function of this gene is only manifested under conditions of mechanical stimulation, and its knockout induces no phenotypic changes related to bone metabolism under sedentary conditions.

### Membrane tension decreases during fusion from pre-osteoclast to mature osteoclasts

Osteoclast maturation is accompanied by increases in cell volume, which may change membrane tension. Therefore, we used a FLIM microscope to investigate the membrane tension of mature osteoclasts (fluorescence lifetime is proportional to membrane tension). First, we defined the different stages of osteoclasts as follows: bone marrow cells stimulated with M-CSF for five days were defined as BMMs; BMMs stimulated with receptor activator of nuclear factor-κB ligand (RANKL) for two days were defined as pre-osteoclasts (pOCs); and BMMs stimulated with RANKL for five days were defined as mature osteoclasts (mOCs) (Figure 2. A). By detecting BMMs, pOCs, and mOCs, we found no significant difference in fluorescence lifetime between BMMs (mean value: 4.87 ns) and pOCs (mean value: 4.88 ns), but the fluorescence lifetime of mOCs (mean value: 4.75 ns) decreased significantly (Figure 2. B–C). These results indicate that membrane tension decreases during the fusion of pOCs into mOCs. We then used RNA sequencing (RNA-seq) data from the GEO database (GSE226625) ^16^ to analyze osteoclasts at different time points; in this sequencing experiment, the time point before RANKL addition was designated as D0, while the second, third, and fourth days after RANKL addition were designated as D2, D3, and D4, respectively, thus forming four groups. (Figure 2. D). Compared with D0, the expression of genes related to mechanotransduction after adding RANKL changed significantly. Mechanical-related signaling pathways were also significantly enriched (Figure 2. E). Notably, D4 is close to our definition of mOC. In the comparison between D4 and D0, we found that mechanical-related signals were gradually enriched in the following two signaling pathways: Cellular response to mechanical stimulus (GO:0071260) and Mechanosensitive monoatomic ion channel activity (GO:0008381). We analyzed the expression patterns of differentially expressed genes (DEGs) in these two signaling pathways, and the corresponding heatmap revealed that the expression levels of most genes decreased at D4 (Figure 2. F). Mechanical stimulation likely affects the fusion and maturation of osteoclasts. Subsequently, we prepared two types of bone sections and measured their stiffness using AFM (Figure 2. G–H). Osteoclasts also formed more actin rings on softer bone slices compared to harder ones (Figure 2. I–J).

**Figure 2.**
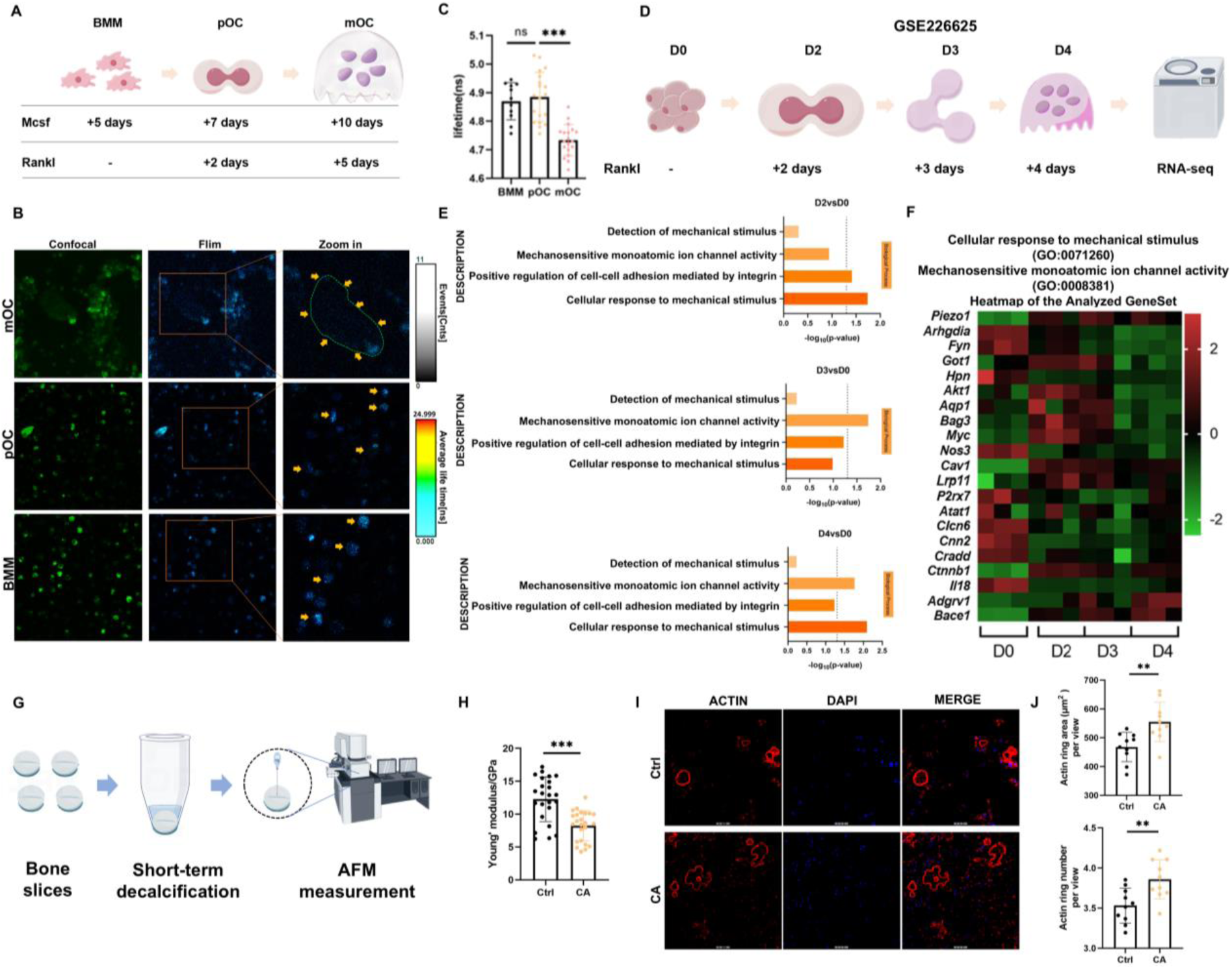
Mechanical signals modulate osteoclast differentiation. **(A)** Schematic of osteoclast culture protocol and osteoclast differentiation process. **(B–C)** Detection of membrane tension of osteoclasts by FLIM microscopy and lifetime statistics (n=18). **(D)** Sequencing grouping schematic (GSE226625) ^16^. **(E)** GO enrichment analysis. **(F)** Heatmap of DEGs. **(G)** Schematic of bone slice processing. **(H)** Stiffness of bone slices measured by AFM (n=24). **(I)** Bone resorption experiments on bone slices. **(J)** Number and area of actin rings (n=10). Data are expressed as the mean ± SD. * P<0.05, * * P<0.01, *** P<0.001.

In summary, membrane tension is downregulated during the fusion of osteoclasts, and mechanotransduction-related genes/pathways undergo stage-specific regulation during RANKL-induced osteoclast differentiation. Furthermore, decreased bone matrix stiffness promotes the formation of core functional structures in osteoclasts, suggesting that bone resorption has a positive feedback property.

### Activation of Piezo1 inhibits maturation from pre-osteoclast to mature osteoclasts

Mechanotransduction relies on mechanically sensitive ion channels (mechanical transducers) to initiate a series of downstream signaling pathways, leading to the regulation of specific cellular and physiological responses ^17^. As one of the most important mechanically sensitive ion channels, Piezo1 mediates cell mitosis, whereas inhibition of Piezo1 leads to cellular multinucleation ^18–20^. Conversely, whether Piezo1 also plays an important role during cell fusion remains to be clarified. We showed that the membrane tension of osteoclasts significant decreased during osteoclast fusion. This suggests that osteoclasts are sensitive to mechanical stimulus and that this process is mediated by Piezo1.

Subsequently, we measured the expression levels of Piezo1 at different stages of osteoclast differentiation (RNA-Seq from GSE226625); its expression increased in the early stage (D0, D2 and D3) but decreased in the later stage (D4) (Figure 3. A). Similarly, we measured the expression levels of *Piezo1* in BMMs, pOCs, and mOCs (which correspond to D0, D2, and D5, respectively) using RT–PCR and found that it decreased significantly in mOCs. (Figure 3. B) and activated by Yoda1 in pOC. TRAP staining and actin fluorescence confirmed that Yoda1 significantly inhibits the fusion and maturation of osteoclasts. (Figure 3. C–D). In addition, Yoda1 inhibited the expression of osteoclast differentiation–related genes (Figure 3. E). Similarly, Yoda1 treatment weakened the bone resorption of osteoclasts on bone slices (Figure 3. F). We overexpressed *Piezo1* on BMMs using transfection reagent and P-ECMV plasmid (S. Figure 1. A–B) to induce BMMs. Consistent with the effects of agonists, overexpression of *Piezo1* inhibited osteoclast differentiation (S. Figure 1. C–G).

**Figure 3.**
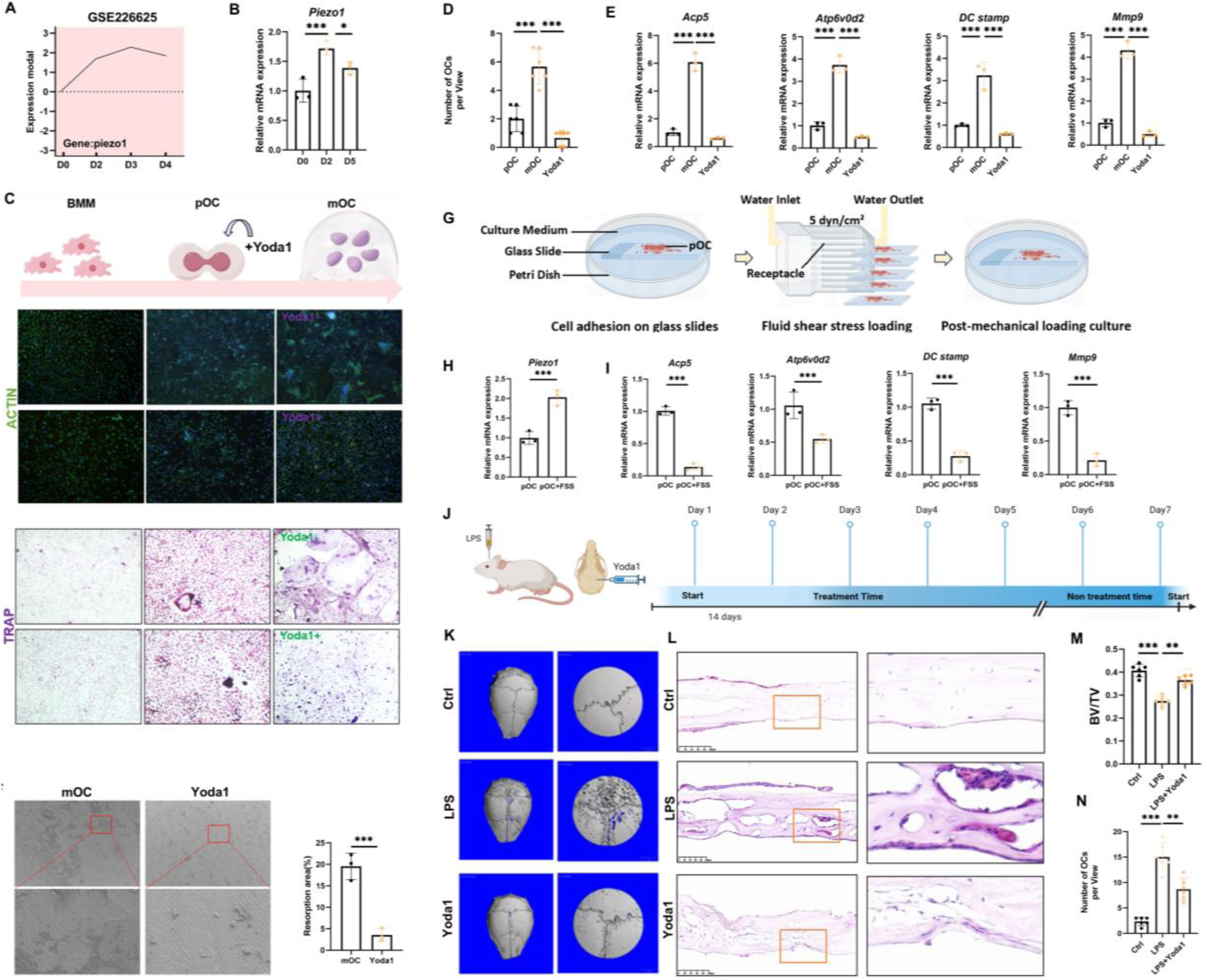
Activation of Piezo1 inhibitors Osteoclast differentiation. **(A)** The expression trend of Piezo1during osteoclast differentiation. **(B)** RT–PCR detection of Piezo1 expression (n=3). **(C)** Actin fluorescence staining and TRAP staining. **(D)** Statistics of osteoclast number (n=6). **(E)** Detection of osteoclast differentiation–related genes (n=3). **(F)** Bone resorption experiment and absorption area statistics (n=3). **(G)** Schematic of fluid shear stress loading. **(H–I)** RT–PCR detection of Piezo1 and osteoclast differentiation–related genes in pOCs **(J)** Schematic of modeling and treatment process. **(K)** 3D reconstruction of mouse skull micro-CT. **(L)** Histological TRAP staining. **(M)** BV/TV statistical analysis (n=6). **(N)** Statistics performed on osteoclast measurements (n=6). Data are expressed as the mean ± SD. * P<0.05, * * P<0.01, n=3. Data are expressed as the mean ± SD. * P<0.05, ** P<0.01, *** P<0.001.

Furthermore, we seeded BMMs onto glass slides and induced their differentiation into pOCs (Figure 3. G). The cells were then placed in a fluid shear stress device to load shear stress, followed by continuous culture; subsequently, we detected Piezo1 and osteoclast differentiation–related genes in pOCs, finding that a shear stress of 5 dyn/cm² could activate Piezo1 and inhibit the differentiation of pOCs into mOCs (Figure 3. H–I).

We isolated BMMs from the bone marrow of two mouse strains, namely WT and *CTSK ^Cre^; Piezo1 ^fl/fl^* and induced their differentiation into mOCs (S. Figure 2. A). Osteoclast-specific Piezo1 knockout did not affect their subsequent fusion (S. Figure 2. B–C). This suggests that Piezo1 does not play a role during the osteoclast fusion process.

LPS can induce osteoclast activation and thereby lead to osteolysis. As shown in Figure 3. J, injection of Yoda1 was performed concurrently with LPS injection into the skull. After 2 weeks, micro-CT showed that Yoda1 treatment significantly alleviated skull osteolysis (Figure 3. K–M). Histological staining revealed that Yoda1 treatment inhibited activation of osteoclasts in the skull (Figure 3. L–N). Thus, Piezo1 does not participate in osteoclast fusion, but it can regulate osteoclast differentiation, maturation, and bone resorptive activity via mechanotransduction. Under pathological conditions, activation of Piezo1 can inhibit aberrant activation of osteoclasts and attenuate pathological osteolysis, which suggests that it exerts a potential protective effect in bone metabolic diseases.

### Piezo1 promotes YAP entry into the nucleus through E-cadherin/Merlin

The YAP signaling pathway has been widely reported as a downstream effector of mechanical stimulation ^21,22^. More importantly, YAP signaling has recently been shown to negatively regulate the differentiation of osteoclasts ^23^. Therefore, we speculate that YAP signaling mediates the effects of Piezo1 activation and the differentiation of osteoclasts. First, we measured the nuclear integration level of YAP in the pOC treated with Yoda1 (Figure 4. A). Yoda1 treatment resulted in shuttling of YAP into the nucleus from cytoplasm (Figure 4. B–C). Similarly, after loading a fluid shear stress of 5 dyn/cm² that can activate Piezo1 (S. Figure 3. A), the nuclear translocation level of YAP in pOCs also increased (S. Figure 3. B–C).

**Figure 4.**
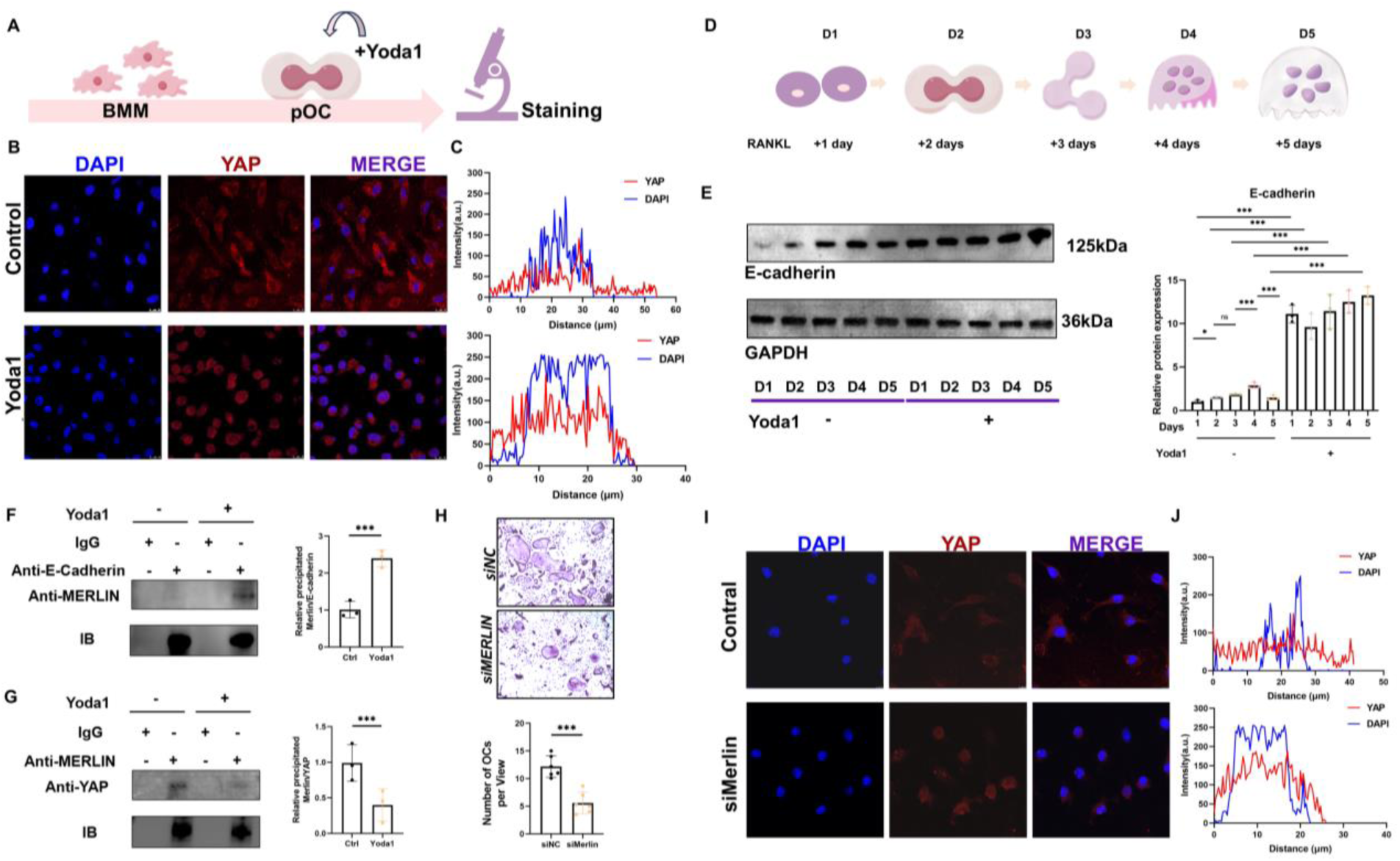
Piezo1 promotes YAP entry into the nucleus through E-cadherin/Merlin. **(A)** Schematic of Yoda1 treatment time points. **(B–C)** Immunofluorescence and quantification of YAP. **(D)** Summary of D0–D5. **(E)** Western blot detection and quantification of E-cadherin (*n*=3). **(F)** COIP detection and quantification of Merlin/E-cadherin (*n*=3). **(G)** COIP detection and quantification of Merlin/YAP (*n*=3). **(H)** TRAP staining and statistics on osteoclast measurements (*n*=6). **(I–J)** Immunofluorescence and quantification of YAP. Data are expressed as mean ± SD. * *P*<0.05, * * *P*<0.01, *** *P*<0.001.

Recent studies have shown that E-cadherin functionally regulates Piezo1 ^12^; E-cadherin is an adhesive molecule that has been reported to bind to Merlin ^24^, an important regulatory molecule of the Hippo pathway, which can bind to YAP and transport it to the nucleus ^24,25^. During the normal differentiation of osteoclasts, E-cadherin typically first increases and then decreases (Figure 4. D–E). It decreases more significantly on the fifth day after the addition of RANKL. However, while activating Piezo1, Yoda1 maintained E-cadherin in a highly activated state. (Figure 4. D–E). We speculate that after E-cadherin is activated, it can anchor Merlin and reduce the binding of Merlin and YAP. Protein immunoprecipitation confirmed that Yoda1 increased the binding of E-cadherin and Merlin, whereas the binding of Merlin and YAP decreased significantly (Figure 4. F–G). To elucidate the role of Merlin, we knocked down Merlin in BMMs and observed its effects on osteoclast differentiation and the nuclear translocation of YAP. Merlin knockdown significantly downregulated the differentiation capacity of osteoclasts and led to increased nuclear translocation of YAP (Figure 4. H–J). In summary, we first clarified the mechanism by which Piezo1 promotes the nuclear translocation of YAP in osteoclasts. Specifically, due to their structural linkage, Piezo1 induces hyperactivation of E-cadherin, which in turn sequesters Merlin—the transport molecule of YAP—resulting in nuclear retention of YAP.

### YAP regulates osteoclast differentiation through NF-κB signaling

We showed that Piezo1 can regulate YAP signaling, but it is unclear whether YAP signaling mediates the effect of Piezo1 on osteoclast differentiation. Subsequently, we used YAP antibodies to perform ChIP-Seq on Yoda1-treated pOC (Figure 5. A) and found that most of the binding regions between YAP and target genes occurred in the promoter region (34.3%) (S. Figure 4. A). We then conducted GO analysis on the enriched target genes, revealing significant enrichment of the NF-κB signaling pathway (Figure 5. B). Previous studies have shown that NF-κB is a key upstream regulator of the expression of NFATc1, the master regulatory switch for osteoclast differentiation ^26^. By consulting the KEGG database, we elucidated all signaling pathways related to osteoclast differentiation (S. Figure 5). The NF-κB signaling pathway was identified at the intersection of GO analysis and the KEGG database. We hypothesized that YAP participates in the NF-κB signaling pathway to regulate osteoclast differentiation and maturation. Therefore, we evaluated the genes related to the NF-κB pathway in the ChIP-Seq. Among 28 genes, 17 genes positively regulate NF-κB signaling and 11 play a negative regulatory role (S. Figure 4. B). Based on the results of the previous GO analysis (GO Term: Negative regulation of NF-κB signaling), we focused on the genes that are negatively regulated (Figure 5. B). Among these 11 genes, we focused on the genes that bind to YAP in the promoter region. Finally, we screened out seven genes. Subsequently, we tested the effects of Yoda1 on these genes. *Tnfaip3* and *Tax1bp1* were significantly upregulated by Yoda1 (Figure 5. C) (S. Figure 6). This suggests that *Tnfaip3* and *Tax1bp1* mediate the mechanical signals transmitted by Piezo1/YAP. In summary, we revealed the specific mechanism of Piezo*1*-promoted nuclear retention of YAP in osteoclasts: Piezo1 is structurally associated with E-cadherin, and its activation induces E-cadherin hyperactivation, which sequesters the YAP transport molecule Merlin, blocks Merlin-mediated YAP nuclear export, and ultimately causes nuclear retention of YAP.

**Figure 5.**
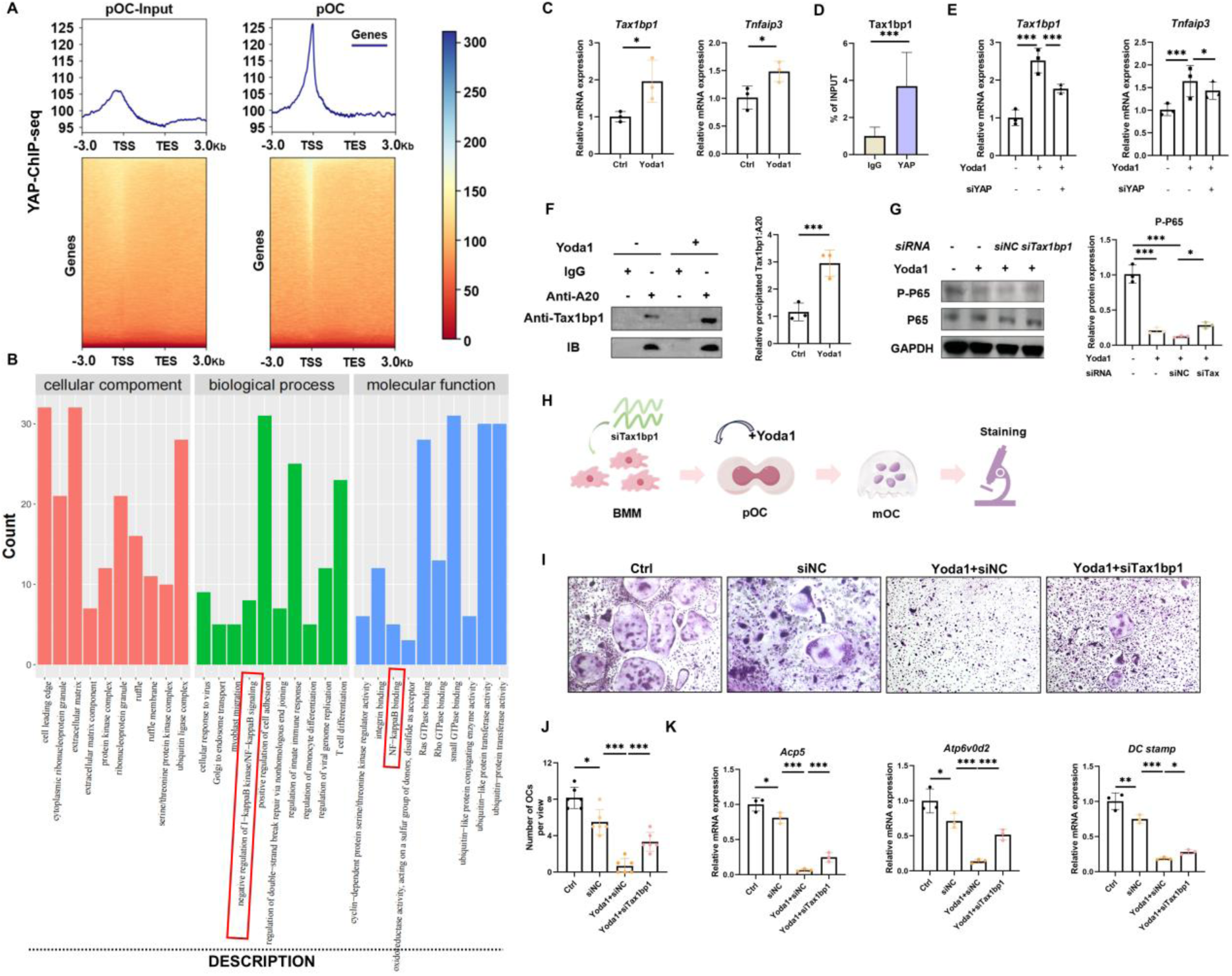
YAP regulates osteoclast differentiation through NF-κB signaling. **(A)** DNA fragment stacking peak at the binding site of YAP on the genome. **(B)** GO analysis of YAP-binding genes. **(C)** RT–PCR analysis of target genes (*n*=3). **(D)** Co-precipitation detection of proteins and DNA. **(E)** RT–PCR detection and quantification of *Tax1bp1* and *Tnfaip3* (*n*=3). **(F)** COIP detection for Tax1bp1 and Tnfaip3 (*n*=3). **(G)** Detection and quantification of phosphorylation levels of P65 (*n*=3). **(H**) Schematic of siRNA transfection and Yoda1 Addition. **(I–J)** TRAP staining and osteoclast count (n=6). **(K)** Detection of osteoclast differentiation–related genes (*n*=3). Data are expressed as the mean ± SD. * *P*<0.05, * * *P*<0.01, *** *P*<0.001

*Tnfaip3* encodes a deubiquitinase called A20, which can form a complex with *Tax1bp1* to negatively regulate NF-κB signaling ^27^. In this process, the main focus is on recruiting and regulating the function of A20 through *Tax1bp1* ^28^. Therefore, we used protein DNA co-precipitation technology to show that Tax1bp1 can bind to YAP antibodies (Figure 5. D). Then, after knocking out *YAP*, we activated Piezo1 using Yoda1 and detected *Tax1bp1* and *Tnfaip3*. We found that in the absence of YAP, Yoda1 was unable to activate *Tax1bp1* and *Tnfaip3* (Figure 5. E). Compared to *Tnfaip3*, *Tax1bp1* was downregulated more significantly in the absence of YAP. Therefore, we speculate that Yoda1 promotes the formation of complexes of *Tnfaip3*- and *Tax1bp1*-encoded proteins and negatively regulates NF-κB signaling. Subsequently, we detected the binding of *Tnfaip3*- and *Tax1bp1*-encoded proteins in Yoda1-treated pOC. Yoda1 promoted the combination of the two (Figure 5. F). By knocking out *Tax1bp1*, we activated Piezo1 with Yoda1 and detected the phosphorylation of P65. In the presence of *Tax1bp1*, Yoda1 can significantly inhibit the phosphorylation of P65; this is weakened in the absence of *Tax1bp1* (Figure 5. G). Furthermore, we knocked down *Tax1bp1* in BMMs, differentiated them into pOCs, and then added Yoda1 (Figure 5. H). Subsequently, the cells were induced to differentiate into mOCs, followed by TRAP staining (Figure 5. I). *Tax1bp1* knockdown rescued the inhibitory effect of Yoda1 on osteoclasts (Figure 5. J–K).

In summary, Yoda1 promotes the formation of a complex between Tax1bp1 and A20—a deubiquitinase encoded by Tnfaip3—via the YAP signaling pathway, thereby inhibiting the activation of the NF-κB signaling pathway (i.e., suppression of P65 phosphorylation), and Tax1bp1 is a key mediator of Yoda1-induced inhibition of osteoclast differentiation.

### Activation of Piezo1 increases bone mass through inhibiting osteoclast activity

During development, mice undergo bone modeling, where the bone marrow cavity will continuously expand under the action of osteoclasts. In the present study, we injected Yoda1 into 4-week-old mice for 2 weeks (Figure 6. A). Yoda1 significantly increased bone mass in mice (Figure 6. B–C). Meanwhile, Yoda1 reduced the inner diameter of the mouse bone marrow cavity (Figure 6. D–E). This indicates that Yoda1 directly inhibits the action of osteoclasts. Dynamic bone histomorphometric analysis following calcein green and tetracycline labeling showed that the bone mineral appearance rate (MAR) of trabecular bone increased after Yoda1 treatment (Figure 6. F–G). Serum test results revealed a decrease in CTX and an increase in PINP following Yoda1 treatment (Figure 6. H). TRAP staining showed a decrease in osteoclasts in the middle section of the tibia after Yoda1 treatment (Figure 6. I–J). These data suggest that Yoda1 directly inhibits osteoclast function while promoting osteogenesis, thereby regulating bone modeling and increasing bone mass in mice.

**Figure 6.**
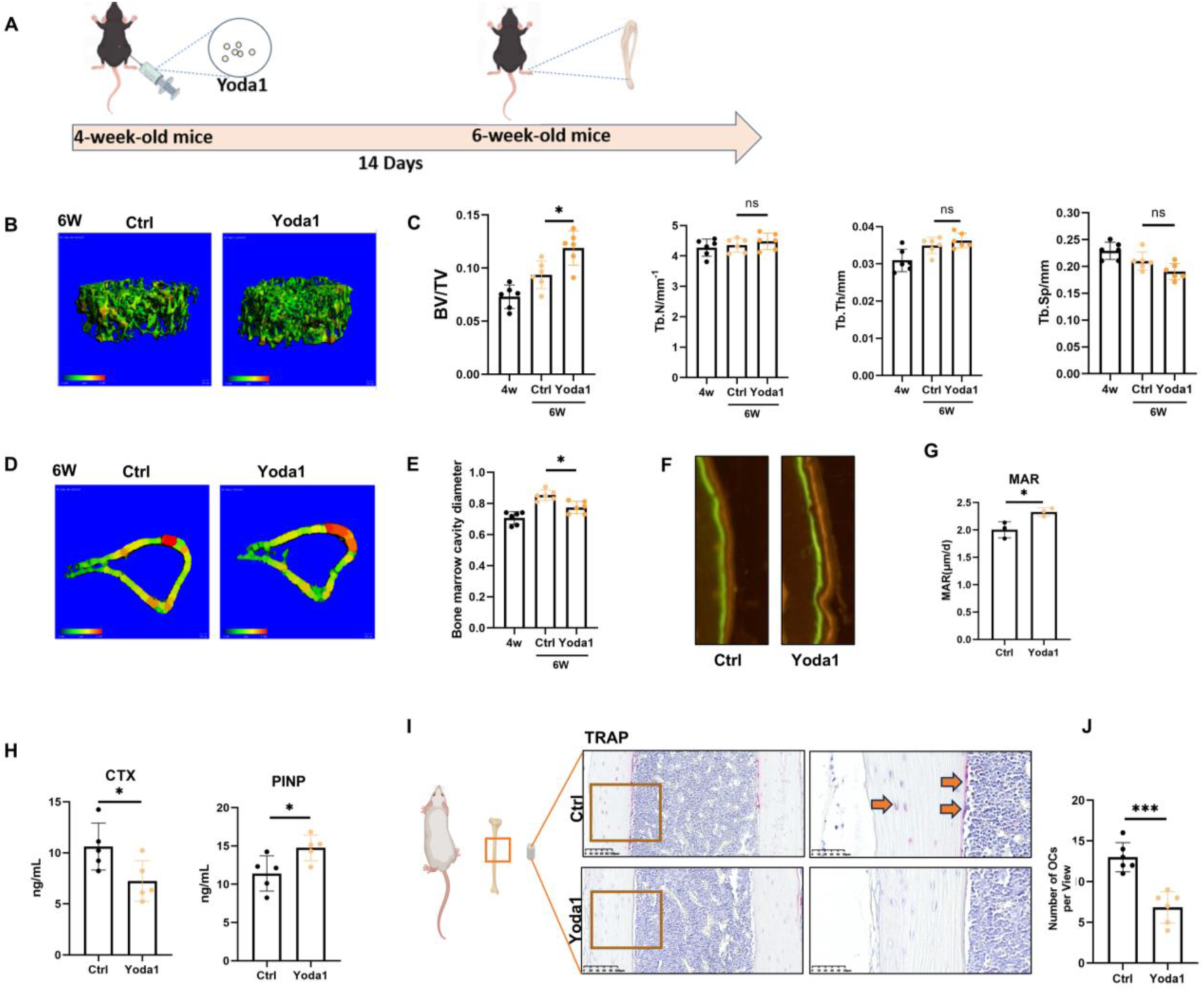
Activation of Piezo1 inhibits osteoclast activity during bone modeling process. **(A)** Schematic of Yoda1 Injection. **(B–C)** µ-CT reconstruction and parameter statistics of tibial trabecula (*n*=6). **(D–E)** Microscopic CT 3D reconstruction of tibial cortical bone and statistical analysis of bone marrow cavity diameter (*n*=6). **(F–G)** Dual labeling of calcein and tetracycline, and MAR statistical analysis (*n*=3). **(H)** ELISA detection of serum CTX and PINP. (*n*=6). **(I–J)** Histological TRAP staining and osteoclast count (*n*=3). Data are expressed as mean ± SD. * *P*<0.05, * * *P*<0.01, \*\*\**P*<0.001

## DISCUSSION

This study identified Piezo1 acts as a core regulatory factor for osteoclasts to sense and transduce mechanical load signals during differentiation. In vitro experiments confirmed that Piezo1 knockout had no effect on the fusion of osteoclast precursors, whereas Piezo1 activation by Yoda1 markedly inhibited this process. This indicates that Piezo1-mediated mechanical stimulation is not essential for normal osteoclast differentiation, yet excessive activation of this pathway disrupts the fusion homeostasis of osteoclasts.

Regulation of mechanical signaling is a major focus in bone biomechanics, and our findings on Piezo1-knockout osteoclasts are consistent with the phenotypic results of previous study ^14^. We further found that both Yoda1-induced Piezo1 activation and increased bone matrix stiffness significantly inhibited the differentiation and fusion of osteoclasts in vitro. Existing studies have reported that osteoclasts primarily rely on cell membrane tension to perceive extracellular mechanical signals, and the Piezo1 channel is an important sensor of membrane tension ^2,6–8,29^. Our results also confirmed that osteoclast fusion is accompanied by a decrease in membrane tension under physiological conditions, which provides a mechanistic explanation for the inhibition of osteoclast fusion by Yoda1.

Previous studies have reported structural interactions between Piezo1 and the key cell adhesion molecule E-cadherin ^12^. Our study further validated this association at the functional level: Piezo1 activation markedly upregulated the E-cadherin signaling pathway in osteoclasts. In addition, E-cadherin is an important regulator of osteoclast differentiation and fusion, and its signaling activity is closely correlated with the intercellular fusion of osteoclast precursors ^13^. However, we confirmed that excessive activation of E-cadherin is detrimental to subsequent osteoclast differentiation.

NF-κB is a pivotal regulatory target for osteoclast differentiation ^30^, and this work verified that Piezo1 activation modulates the expression and activation of NF-κB via the YAP signaling pathway. Although Piezo1-mediated regulation of YAP has been widely reported in various cell types, the specific molecular mechanism underlying this regulatory process in osteoclasts remains elusive ^22,25,31^. Our study for the first time uncovers a regulatory cascade in osteoclasts: Piezo1 activation promotes the binding of E-cadherin to Merlin and reduces Merlin–YAP interaction. As a key regulator of YAP, Merlin can induce the nuclear translocation and degradation of YAP, and altered Merlin binding status is a core step leading to YAP signaling inactivation. This finding fills the research gap in the specific mechanism by which Piezo1 modulates YAP in osteoclasts and further elucidates the molecular basis underlying the inhibition of osteoclast differentiation by Piezo1 activation via the NF-κB pathway (Figure 7).

**Figure 7.**
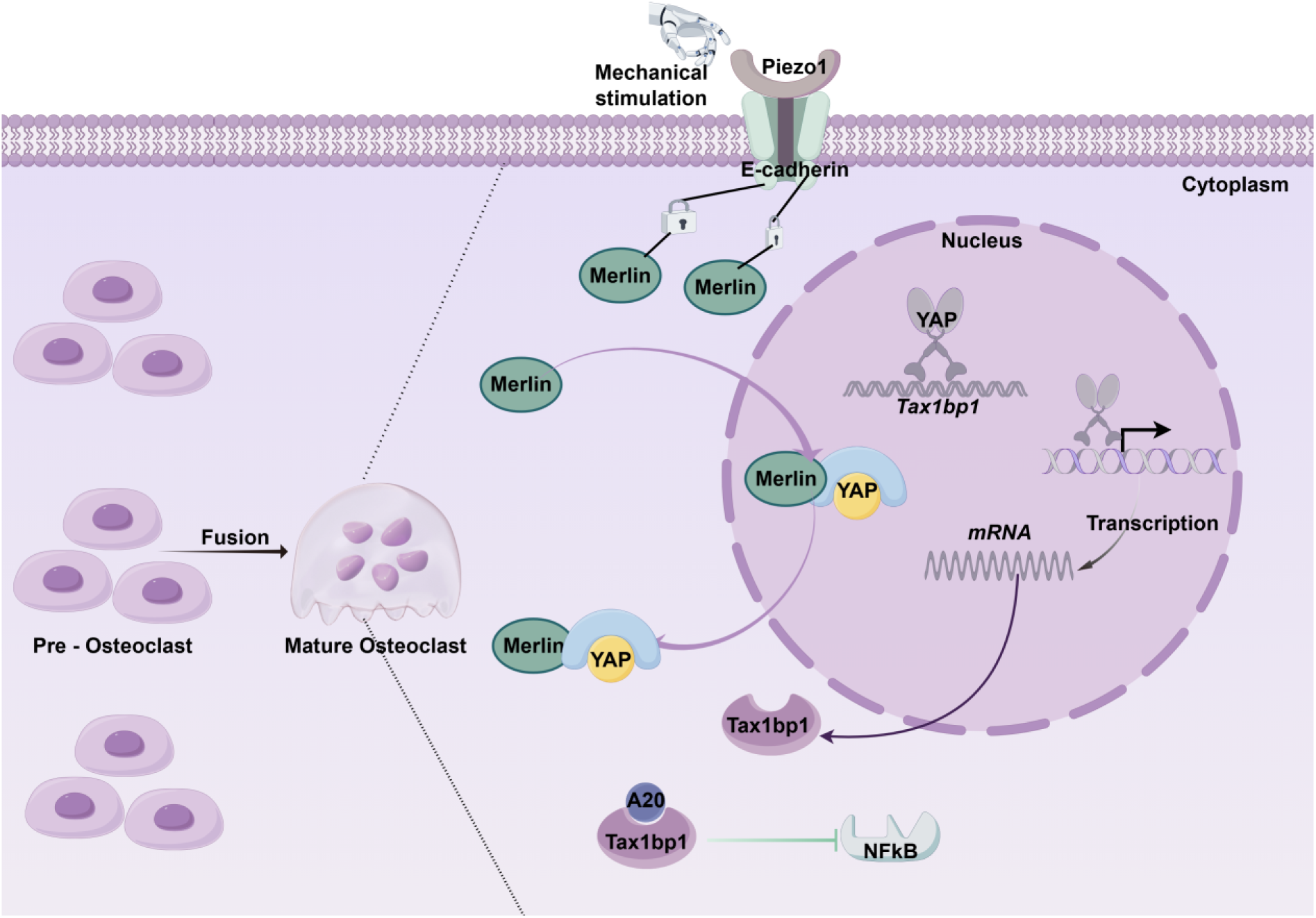
Piezo1-mediated mechanical signaling inhibits osteoclast fusion and regulates bone homeostasis via the E-cadherin–Merlin–YAP–NF-κB pathway.

However, our study has several limitations. First, in vivo validation of key effector molecules such as *Tax1bp1*, particularly the generation of gene knockout mice, was not performed and would help substantiate our conclusions. Second, we did not verify our findings with clinical samples; specifically, the expression levels of key molecules including Piezo1, E-cadherin, and Tax1bp1 in bone tissues of patients with weightlessness-induced osteoporosis (e.g., astronauts) and disuse osteoporosis remain to be measured to establish the correlation between the conclusions from animal experiments and human physiology. Finally, an important conclusion throughout our study is that osteoclast-specific Piezo1 knockout does not result in obvious bone phenotypes in mice. However, this cannot rule out the potential role of compensatory mechanisms. For instance, in studies on parturition mechanics, specific Piezo1 knockout alone in the uterus or sensory neurons causes only mild parturition abnormalities, as Piezo2 compensates for some functions of Piezo1. Only concurrent knockout of both Piezo1 and Piezo2 significantly impairs normal parturition ^32^.

The skeletal system provides a mechanical framework for the entire body. The reduction in bone mass caused by aging or hormone-related osteoporosis can be treated through different strategies, including synthetic metabolism therapy that increases bone formation and catabolic metabolism therapy that reduces bone resorption ^33^. In fact, astronauts typically lose more bone mass within a month than postmenopausal women on Earth within a year, and the decrease in bone strength is even greater ^34^. However, bone loss caused by microgravity or lack of mechanical force cannot be effectively suppressed, indicating that the molecular mechanism of bone loss caused by lack of mechanical force may be different from that of aging or hormone-related osteoporosis. Our study suggests that Piezo1 can regulate differentiation fate by directly sensing the mechanical load of osteoclasts. Our findings will help guide the development of novel anti-bone resorption treatments of osteoporosis.

## MATERIALS AND METHODS

### Animals

All male C57BL/6 mice were purchased from Shanghai Jihui Laboratory Animal Care Co., Ltd. (Shanghai, China). For the treadmill exercise, the 8-week-old mice underwent a progressive acclimatization period over 2 days, starting with a low incline and slow speed to familiarize them with the exercise equipment ^35^. Over the next 2 days, the incline was moderately increased, and the exercise duration was extended gradually. From day 5 onward, the incline was further increased, and the speed was fine-tuned according to the mouse tolerance. After acclimatization, all exercise sessions were performed at a speed of 24 m/min and an incline of 5°, with each session lasting 1 h. The chronic training protocol consisted of daily exercise for 28 consecutive days. Immediately after the final exercise session, blood samples were collected from the mice, followed by tissue harvesting.

Yoda1 was administered in 4-week-old mice via intraperitoneal injection on the first 5 days of each week, with no injections given on the remaining 2 days ^19^. After 2 consecutive weeks of this injection regimen, tissues were collected. Yoda1 (Sigma-Aldrich, St. Louis, MO, USA) was dissolved in dimethyl sulfoxide at a concentration of 40 mM to prepare the stock solution. After dilution with 5% ethanol, the solution was administered to mice via intraperitoneal injection at a dose of 5 µmol per kg of body weight. Mice were sorted by body weight and then assigned to either the vehicle control group or Yoda1 group, ensuring that the average body weight was identical across all groups.

For the osteolysis model, a blade was used to cut open the skin on the top of the 8-week-old mouse head and scrape off the periosteum prior to injection of lipopolysaccharide (LPS) (200 g/mouse in a 100-μL volume). The injection was administered the first 5 days of each week and not the last 2 days, with a total cycle of 2 weeks. For treatment of the bone resorption model, Yoda1 (5 μmol per kg) was concomitantly administered at the same site as the LPS injection. The cpbsontrol group received the same volume of phosphate-buffered saline (PBS). Samples were collected 2 weeks later.

All researchers who collected data were blinded to the mouse genotypes and their respective group assignments. This study received ethical approval, and all animals used in this study received care in accordance with the institutional guidelines developed by the Animal Experiment Ethics Committee of Shanghai Jiao Tong University School of Medicine (No. SH9H-2023-A3-1).

### Micro-computed tomography (μCT) analysis

Right tibias were scanned using a high-resolution μCT scanner (μCT 80; Scanco, Zurich, Switzerland) to obtain the trabecular and cortical bone microstructure (BV/TV, Tb.N, Tb.Sp, Conn.D, Ct.Th, and SMI). The scanning parameters were set as follows: voltage, 70 kV; electric current, 114 μA; and resolution, 10 μm per voxel. For trabecular measurements, a region of interest (ROI) was defined as 1.9 mm from the proximal tibial condyles, immediately distal to the growth plate, and extended to 100 slices. For cortical bone analyses, an ROI was defined at the mid-diaphysis, starting 4.5 mm from the proximal tibial condyles and extending to 100 slices.

For skull analysis, the ROI was manually delineated as the bilateral parietal bone regions selected symmetrically along the skull midline. Three-dimensional–reconstructed models of the skull were generated using software (SkyScanNRecon) to visually display the location and extent of osteolytic regions. Raw projection images, reconstructed volume data, ROI annotation files, and parameter analysis were categorized and stored according to the “sample number - scan date” format.

### Atomic force microscopy (AFM) measurement

The prepared bone slices were fixed on a glass slide for AFM (TPP Technology Plastic Products Company, Trasingen, Switzerland), using a biocompatible sample adhesive (JPK Instruments Company, Berlin, Germany). The AFM system used was a combination of CellHesion 200 (Burker Berlin, Germany) and an inverted phase contrast microscope (AxioObserver D1, Jena Carl Zeiss microscope, Germany). The cantilever used for micrometer scale indentation was calibrated with a spherical tip with a radius of 5 microns (model: SAA-SPH-5UM, k=0.2 N/m, Burker, Billerica, Massachusetts, USA). The linearly fitted area on the calibration force–distance curve was used for calibration and calculation of Young’s modulus.

### Immunohistochemistry stain

Bone samples were immersed in 4% paraformaldehyde (Cat#: P0099, Beyotime) at 4°C for 24 h and decalcified in 10% Ethylene Diamine Tetraacetic Acid (EDTA) (Cat#: BL616A, Biosharp) at 4°C for 4 weeks and then embedded in paraffin. Fixed, paraffin-embedded bone tissue sections (5 μm) were assessed using immunohistochemical (IHC) staining. The sections were deparaffinized and hydrated with distilled water, followed by antigen recovery using 0.25% trypsin. Endogenous peroxidase activity was blocked using hydrogen peroxide. Sections were incubated with primary antibodies overnight at 4°C, followed by incubation with horseradish peroxidase–conjugated secondary antibodies. Positivity rates in the stained areas were calculated using Image-Pro Plus software (Media Cybernetics, Rockville, MD, USA). To quantify bone formation in the periosteum and endosteum of cortical bone, mice were injected with calcein (20 mg/kg body weight) and alizarin red (20 mg/kg body weight) at 10 and 3 days prior to tissue collection, respectively.

### Cell culture

To cultivate bone marrow–derived macrophages (BMMs), bone marrow cells were extracted and immersed in MEM-α culture medium containing fresh 10% fetal bovine serum with 25 ng/mL Macrophage Colony-Stimulating Factor (M-CSF). Culture medium was refreshed every 2–3 days. BMMs were then induced into osteoclasts using a culture medium containing 25 ng/mL M-CSF (Cat #: RP01216, abclonal) +50 ng/mL Rankl (Cat#:462-TEC-010, R&D Systems). After 5 days, the stimulation was terminated by observing the fusion of osteoclasts. After fixing the cells, they were stained with TRAP solution according to the instructions of the Sigma TRAP staining kit. After co-culturing at 37°C in the dark for 30–45 minutes, images were acquired (Leica DMi8, Germany).

### RNA extraction and reverse transcription–quantitative polymerase chain reaction (RT–qPCR)

Total RNA was extracted from the cultured cells using the AxyPrep Multisource RNA Miniprep Kit (Cat#: AP-MN MS-RNA-250, Axygen, Corning, New York, USA). Briefly, after the cells were lysed, centrifugation and washing were performed using various reagents from the kit according to the manufacturer’s instructions. Finally, TE buffer was added to dissolve the RNA, which was then centrifuged to obtain a liquid RNA sample. Complementary DNA (cDNA) was synthesized via reverse transcription using TaKaRa Reverse Transcription Reagent (Cat#: D2680A, TaKaRa, Japan). RT–qPCR was performed using a QuantStudio 6 Flex RT qPCR System (Applied Biosystems, CA, USA) and SYBR Green PCR Mix (Cat#: B21402, Bimake, TX, USA). The relative RNA level was calculated using the comparative threshold cycle (2^−ΔΔCT^) method and normalized to the value of Gapdh within the sample.

### Western blot analysis

For the extraction of total proteins, cells were lysed for 15 min using cell lysis buffer (Cat#: P0013C, Beyotime, China) containing protease inhibitors (Cat#: P1005, Beyotime, China) and then sonicated. The collected protein solution was mixed with sampling buffer (Cat#: P0015, Beyotime, China) and incubated at 99°C for 10 minutes. Proteins were then separated with 4%–20% ExpressPlus™ PAGE Gel (Cat#:M01215C, GenScript) and electrophoresed in Tris-MOPS-SDS Running Buffer (Cat#:M00677, GenScript) diluted with ddH_2_O. The gels were then electroblotted onto 0.22-µm PVDF membranes (Cat#: L00735, GenScript). Gels were blocked with 5% BSA–TBST (Tris-buffered saline (TBS)–0.1% Tween 20) (Cat#:ST673, Beyotime, China) for 1 h at room temperature. The membrane was incubated with primary antibodies overnight at 4°C. The next day, the membranes were washed with PBS and then incubated with an appropriate secondary antibody (Cat#: SA00001-1 or SA00001-2, Proteintech, 1:5000) in 2% non-fat milk for 1 h. The immunoreactive bands were detected using a chemiluminescence kit (Cat#: RPN2232, Amersham Biosciences Ltd., UK) and analyzed using Image-Pro Plus software (version 6.0). The relative protein expression level was normalized to the intensity of the Gapdh band.

### Co-immunoprecipitation

Cells were washed twice with PBS before adding precooled RIPA buffer (1 mL/10^7^ cells); they were then scraped from the culture dish or bottle using a cell scraper, transferred to a 1.5EP tube, and shaken slowly at 4°C for 15 minutes. The cells were centrifuged at 14,000 ×*g* for 15 minutes at 4°C, and the supernatant was immediately transferred to a new tube. Protein A agarose was then prepared by washing the beads twice with PBS and diluting in PBS to a concentration of 50% before shaking at 4°C for 10 minutes to remove non-specific impurities and reduce background. This was followed by centrifuging the mixture at 4°C and 14,000 ×*g* for 15 minutes, transferring the supernatant to a new tube, and removing Protein A beads. The total protein was diluted in PBS to approximately 1 μg/μL. To reduce the concentration of descaling agent in the cracking solution, a certain volume of rabbit antibody was added to 500 μL of total protein. The antigen–antibody mixture was slowly shaken at 4°C overnight. A total of 100 μL Protein A agarose beads were used to capture antigen–antibody complexes, and the antigen–antibody mixture was slowly shaken overnight at 4℃ or at room temperature for 1 h. The mixture was centrifuged at 14000rpm for 5 seconds before collecting the agarose bead antigen–antibody complex, removing the supernatant, and washing three times with precooled RIPA buffer. The agarose bead antigen–antibody complex was suspended in 60 μL of 2× loading buffer and mixed gently. The sample was boiled for 5 minutes and centrifuged with free antigens, antibodies, and beads. The supernatant was electrophoresed, and the remaining agarose beads were collected for electrophoresis.

### Fluorescence lifetime imaging microscopy

Fluorescence lifetime imaging microscopy (FLIM) was performed using the Flipper-TR probe (AmyJet, Cat. No.: CY-SC020) as follows: A 1 mM stock solution of Flipper-TR was prepared and stored as described by the manufacturer. A working solution was prepared from the 1 mM stock solution to stain cultured cells and added to the cell culture medium to a final concentration of 1 μM, followed by replacing the original medium with this staining solution. The cells were returned to a humidified incubator at 37°C with 5% CO₂ and incubated for 15 minutes before proceeding with imaging. Cells were protected from light at all times. The stained cells were imaged using the standard FLIM technique: excitation was performed with a 485 or 488 nm pulsed laser, and photons were collected through a 600/50 nm bandpass filter. First, the cells were located using a confocal microscope before switching to the FLIM microscope for imaging. After image acquisition, cells were manually outlined to calculate their lifetime.

### Chromatin immunoprecipitation sequencing

Cells in the logarithmic growth phase were selected with a quantity of ≥1×10⁷ cells and washed twice with pre-chilled PBS to remove residual culture medium. Pre-chilled 1% formaldehyde solution was added to the collected cell pellet before incubation at room temperature (20–25 °C) for 10–15 minutes. Stop solution (e.g., 2.5 M glycine, with a final concentration of 0.125 M) was then added, followed by incubation at room temperature for 5 minutes to terminate the cross-linking reaction. The mixture was centrifuged at 1000 ×*g* at 4°C for 5 minutes, and then the cell pellet was collected and washed twice with pre-chilled PBS (containing protease inhibitors) to remove residual formaldehyde.

Nuclear lysis buffer was added to the cross-linked cell pellet (approximately 1 mL for 1×10⁷ cells), and lysis was conducted on ice for 10 minutes. The lysate was transferred to sonication-specific centrifuge tubes (avoiding bubbles), and sonication was performed using a sonicator as follows: power 30%–40%, sonication time 30 seconds, interval 30 seconds, for a total of 10–15 cycles. This was followed by centrifugation at 12,000 ×*g* at 4℃ for 15 minutes, collection of the supernatant, measurement of DNA concentration using a Nanodrop, and verification of the fragment size via 1.5% agarose gel electrophoresis (target fragments should be concentrated in the 200–500-bp range).

Anti-YAP antibody was used to capture the target protein, thereby enriching the DNA fragments bound to it. DNA was dissociated and purified from the “bead complex” to remove impurities such as proteins and antibodies. End repair, adapter ligation, and amplification were performed on the purified DNA fragments to construct a library compatible with the Illumina sequencing platform. Subsequent paired-end sequencing (PE150) was conducted.

### Statistical analysis

Data are shown as mean ± standard deviation (SD), and the results are represented as bar graphs with individual data points. The color intensity of the images was measured using ImageJ v1.8.0 software (National Institutes of Health, USA). Statistical analyses were performed using GraphPad Prism Version 9 software (La Jolla, California) and SPSS22.0 (IBM, New York, NY). All data were tested for normality using the Shapiro-Wilk test. An unpaired two-tailed Student’s t-test was used for pairwise comparisons between two groups. For comparisons involving three or more groups, a one-way analysis of variance followed by Tukey’s post-hoc test was used. The non-parametric Mann–Whitney U test was used to compare non-parametric datasets (non-normal distribution or *n*<6) between two groups. *P* ≤ 0.05 was considered statistically significant.

## Author contributions

Conceptualization: Z.Y., H.L., and L.C. Methodology: Y.W., and H.W. Investigation: Y.W., H.W., X.L., T.K., and J.C. Writing—original draft: Y.W., and H.W. Writing—review and editing: Z.Y., H.L., and L.C. Funding acquisition: Z.Y. Resources: K.Y., L.W., and M.Y. Supervision: K.Y., L.W., and M.Y.

## Data and materials availability

All data and reagents necessary for the evaluation and reproduction of the results reported in this paper are contained in the article and/or its supplementary materials. Newly generated materials produced in this study are available upon request by contacting the corresponding author Z.Y. at. All transcriptomic data used in this study have been deposited in the Gene Expression Omnibus (GEO) database of the National Center for Biotechnology Information (NCBI). The accession number for the RNA-sequencing data is GSE226625.

