## Supplementary Figure for "Piezo1 transduces mechanical signals to inhibit osteoclast fusion and coordinate bone homeostasis"

Yao Wang et al.

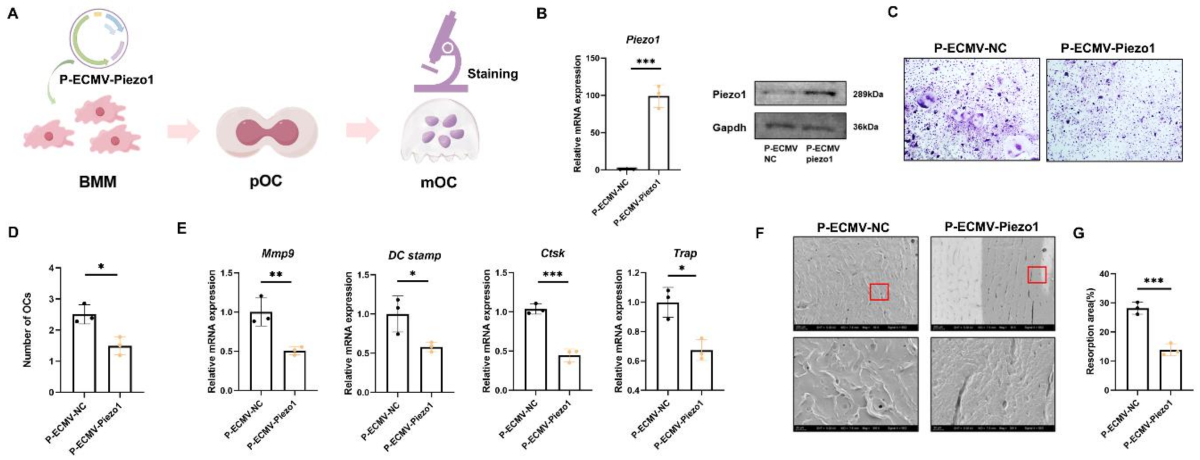


**Supplementary Figure 1: Overexpression of Piezo1 inhibits osteoclast differentiation and maturation. (A–B)** Measurement of *Piezo1* expression levels after overexpression. **(C–D)** TRAP staining and osteoclast count. **(E)** Detection of osteoclast differentiation–related genes. **(F–G)** Bone resorption experiment and absorption area statistics. Data are expressed as mean ± SD. * $P<0.05$, ** $P<0.01$, *** $P<0.001$, $n=3$.

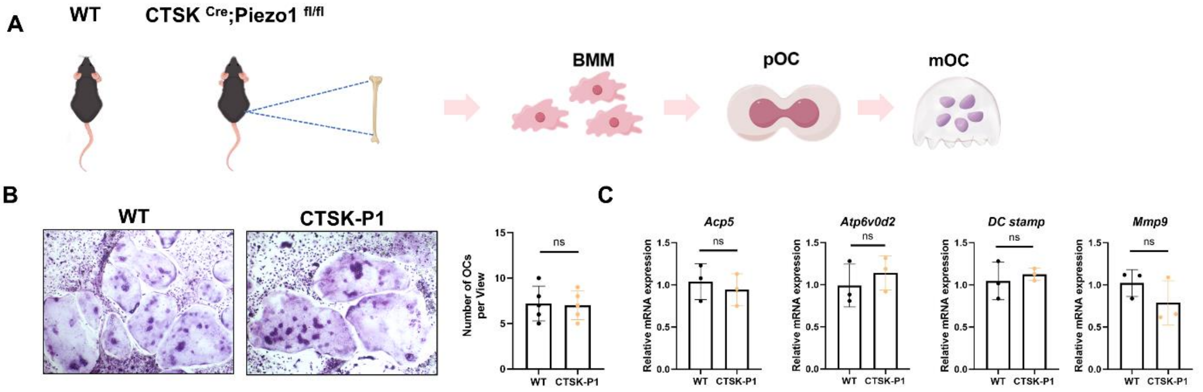


**Supplementary Figure 2: Piezo1 knockout does not affect osteoclast differentiation and maturation. (A)** Schematic of BMM isolation. **(B)** TRAP staining and osteoclast count (*n*=5). **(C)** Detection of osteoclast differentiation–related genes. Data are expressed as mean ± SD. * $P<0.05$, ** $P<0.01$, *** $P<0.001$, $n=3$.

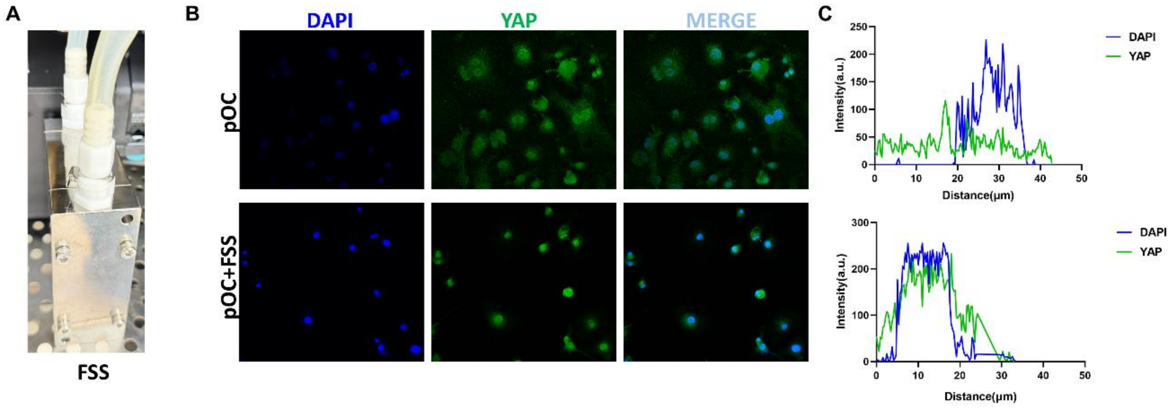


**Supplementary Figure 3: Fluid shear stress application promotes the nuclear translocation of YAP in pOCs. (A)** Physical diagram of the shear module. **(B–C)** Immunofluorescence and quantification of YAP. The data are expressed as the mean ± SD. * $P<0.05$, ** $P<0.01$, *** $P<0.001$, $n=3$.

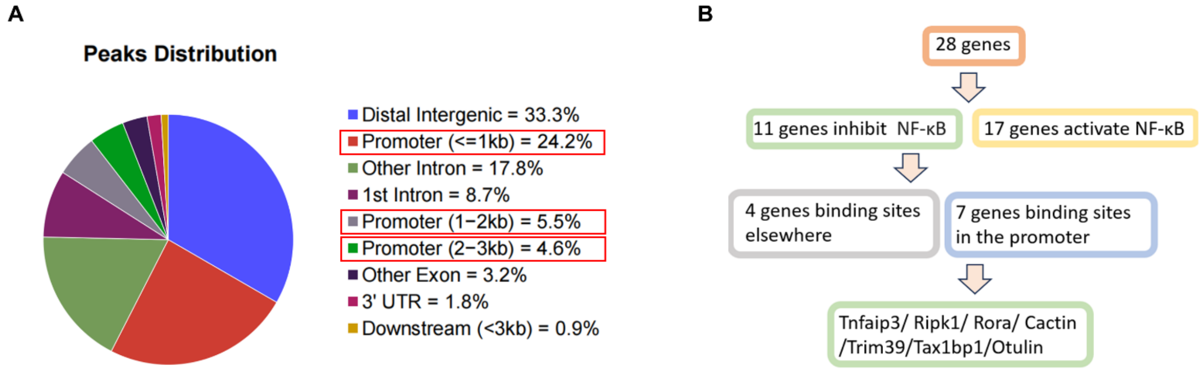


**Supplementary Figure 4: Screening protocol for target genes. (A)** Binding position of YAP on the genome. **(B)** Schematic of target gene screening.

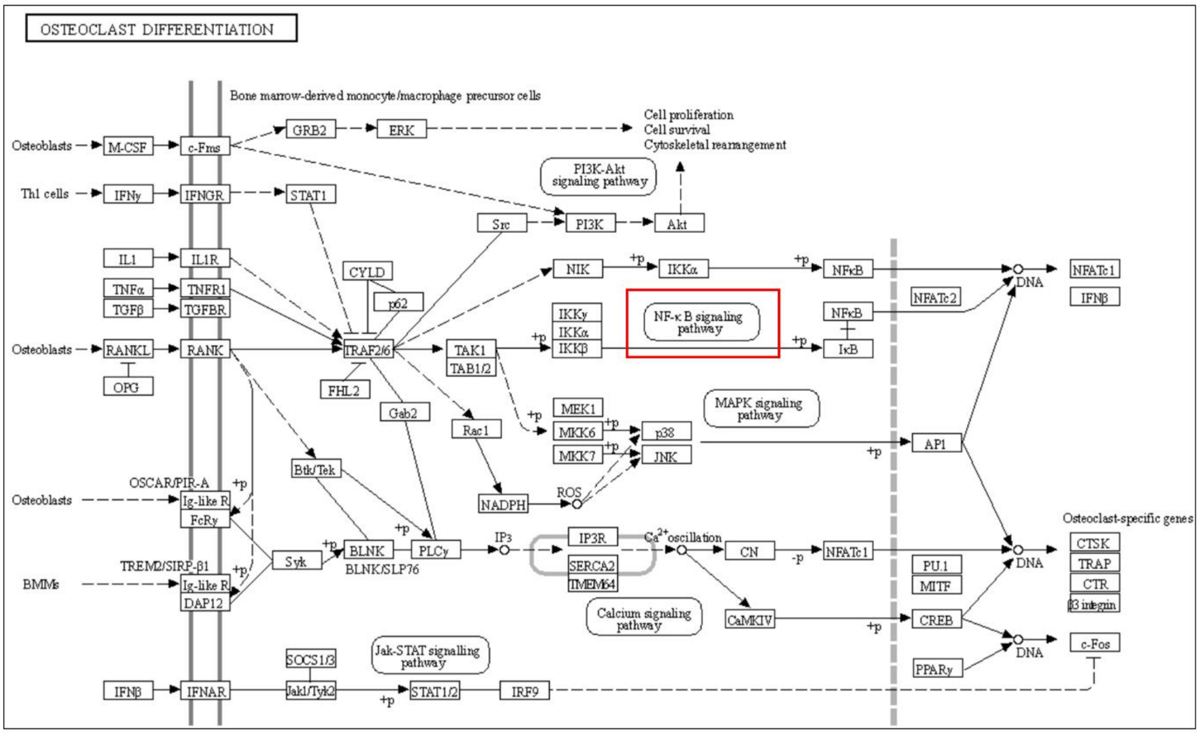


**Supplementary Figure 5: The osteoclast differentiation signaling pathway in the KEGG database.**

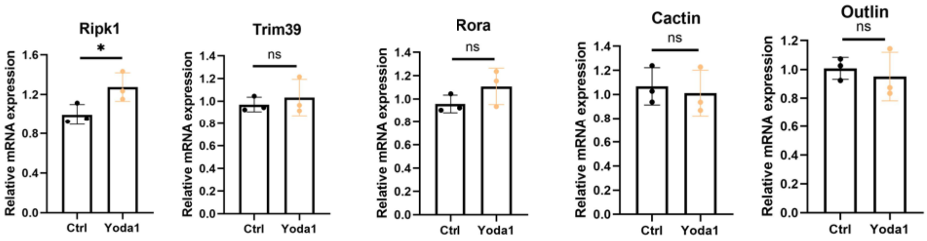


**Supplementary Figure 6: RT–PCR detection of genes from GO term (Negative regulation of NF-κB signaling)**. Data are expressed as mean ± SD. * $P$<0.05, ** $P$<0.01, *** $P$<0.001, $n$=3.
